# Reduced glutathione decreases cell adhesion and increases cell volume

**DOI:** 10.1101/2021.07.30.454460

**Authors:** Alain Géloën, Emmanuelle Berger

## Abstract

Glutathione is the most abundant thiol in animal cells. Reduced glutathione (GSH) is a major intracellular antioxidant neutralizing free radicals and detoxifying electrophiles. It plays important roles in many cellular processes, including cell differentiation, proliferation, and apoptosis. In the present study we demonstrate that extracellular concentration of reduced glutathione markedly increases cell volume within few hours, in a dose-response manner. Pre-incubation of cells with BSO, the inhibitor of γ-glutamylcysteine synthetase, responsible for the first step in intracellular glutathione synthesis did not change the effect of reduced glutathione on cell volume suggesting a mechanism limited to the interaction of extracellular reduced glutathione on cell membrane. Similarly, inhibition of γ-glutamylcyclotransferase involved in intracellular glutamate production had no effect on the action of reduced glutathione. Oxidized glutathione exerted no effect on cell volume. Results show that reduced GSH decreases cell adhesion resulting in an increased cell volume. Since many cell types are able to export GSH, the present results suggest that this could be a fundamental self-regulation of cell volume, giving the cells a self-control on their adhesion proteins.

## 1. Introduction

Oxidative stress results from an imbalance between reactive oxygen species production (ROS) and the abilities of biological systems to eliminate them. In that context, reduced glutathione (GSH) is the central redox agent in most aerobic organisms and a fundamental player in the fine regulation of oxidative stress. GSH is involved in numerous vital functions such as detoxifying electrophiles, scavenging free radicals, maintaining the thiol status of proteins, providing cysteine, modulating DNA synthesis, microtubular-related processes, nitric oxide homeostasis, the activity of neuromodulators as well as immune functions (reviewed in [1]). As a result, reduced GSH plays important roles in cell metabolism, differentiation, proliferation and apoptosis. Disturbances in GSH homeostasis are implicated in numerous human diseases, including cancer, pathologies associated to aging, cardiovascular, inflammatory, immune degenerative diseases [2].

Although GSH is present in the extracellular fluid, it cannot be directly uptaken by cells [1, 2]. It is degraded by cells expressing the ectoprotein γ-glutamyl transpeptidase to produce dipeptides and free amino-acids uptaken by cells and used for *de novo* GSH synthesis. GSH is a tripeptide (γ-L-glutamyl-L-cysteinylglycine) synthesized by two cytosolic ATP dependent enzymes: glutamate cysteine ligase (GCL) leading to the production of γ-glutamyl-cysteine from glutamate and cysteine and glutathione synthetase (GS) involved in the ligation of γ-glutamyl-cysteine to glycine to produce γ-glutamylcysteinylglycine. After its synthesis, almost 90 % is retained into the cytoplasm [2, 3].

Cellular redox regulation are crucial in cell-matrix interaction (4). Adhesion between cells and extracellular matrix is under the control of integrins. They have been identified as a target of redox-regulation by ROS (5). Indeed all integrins contain a large number of cysteine residues in their ectodomains. The protein disulfide isomerase (PDI) regulates integrin-mediated activities, including adhesion. Blocking PDI inhibits adhesion (4, 6, 7). Redox agents such as glutathione were shown to regulate the function of integrins (8, 9). Integrins connect the extracellular matrix to the actin cytoskeleton. It has been suggested that integrins serve as cell volume sensors (10, 11). Despite the described effects of redox on integrins, the effets of reduced glutathione on cell volume has never been reported so far. The presents study relates experimental results pointing out the effect of reduced glutathione on cell volume, using different methods. Implications of such a new effect of reduced glutathione will be discussed in the light of recent data from the litterature.

## 2. Materials and Methods

### 2.1 Cell culture

Human lung carcinoma A549 cells, were grown in Dulbecco’s Modified Eagle’s Medium with low glucose (1g/L) containing 10% fetal calf serum (DMEM 10% FCS, PAA Laboratories, Canada), streptomycin plus penicillin (100 units/mL; Sigma Aldrich, St Quentin-Fallavier, France). Before experiment, cells were dissociated by trypsin (0.05%, Sigma Aldrich) and cell concentration measured using a Sceptor pipette (Millipore, Burlington, USA). Cells were seeded at 5000 cells/cm^2^ in either E-96 plates or in 96 well plates..

### 2.2 Chemicals

Reduced glutathione, buthionine sulfoximine (BSO) and TRITC-phalloïdin were purchased from Sigma-Aldrich. BSO is an effective and specific inhibitor of γ-glutamylcysteine synthetase, the enzyme of the first step in glutathione synthesis [12]. Zombie Red was purchased from Ozyme (Saint-Cyr L’Ecole, France). Intracellular concentration of glutathione was measured using the GSH-Glo(tm) Glutathione Assay from Promega. Pro-GA (Funakoshi Co., Ltd) is a new, pro-drug type inhibitor of γ-glutamylcyclotransferase (GGCT, (EC 2.3.2.4), is part of the gamma-glutamyl cycle, which catalyzes the degradtion of (5-L-glutamyl)-L-amino acid into 5-oxoproline + L-amino acid. 5-oxoproline is a precursor of glutamate, essential for intracellular production of glutathione.

### 2.3 Measurements of cytotoxicity

#### 2.3.1 Real-time cell analysis of impedance

Real-time cell analysis xCELLigence biosensor (RTCA, Agilent, Santa Clara, USA) measures cell surface occupancy (i.e. by impedance measurement), expressed as cell index. It takes into account cell number, cell size and adhesion force. Cells were grown until cell index reached around one (i.e. linear proliferative phase), which may take between 48 to 72 hours. Then culture medium was replaced by a fresh one containing either reduced glutathione and/or different inhibitors. The impedance was recorded every 15 min in a standard 37°C cell culture incubator with 5% CO_2_.

#### 2.3.2 Quantitative phase imaging

Quantitative phase imaging was performed using the Holomonitor M4 digital holographic cytometer (DHC) from Phase Holographic Imaging (PHI, Lund, Sweden). The microscope was housed in a standard 37°C cell culture incubator with 5% CO_2_. Average optical cell volumes were measured in real-time every ten minutes during ten hours in control and after GSH addition.

#### 2.3.3 Cytometry

For cell toxicity assay, at the end of experiments, living cells were incubated during 15 min. at 37°C in the presence of Zombie Red (1/1000) according to the manufacturer’s procedure (Ozyme) then trypsinized and fixed in PBS with formalin 3% (Sigma Aldrich). For F-actin quantification, at the end of experiment, cells were trypsinized then fixed with PBS/ formalin 3% and incubated with PBS/ Triton 0.1%/ TRITC-phalloïdin 1 µg/mL (Sigma Aldrich) during 5 min. Cell count and fluorescence analyses were performed onto a Novocyte cytometer (ACEA) on at least 4 replicates.

#### 2.3.4 Measure of oxidative stress

After treatment cells were trypsinized and fixed with formalin 10% (Sigma, Saint-Quentin Fallavier, France). After 20 min incubation at 4 °C, cells were centrifuged at 1000 rpm for 5 min (Eppendor 5910R, Hambourg, Germany). The pellet was dissolved in 100 µL of dihydrorhodamine 123 (DHR) 1x diluted in PBS Triton 0.1%. The cells were incubated with the DHR solution for a minimum of 15 min at room temperature in the dark. After incubation, cells were analyzed with FACS (Novocyte flow cytometer, Acea Biosciences, San Diego, USA). Data were collected using the Software NovoExpress (Acea Biosciences, San Diego, USA) and manually analyzed using Excel.

### 2.4. Statistical analysis

Data presented are representative experiments performed at least in triplicates, as mean values ± SEM. Statistical analysis was performed with Stat View 4.5 software (Abacus Corporation, Baltimore, USA) for Windows, the data were analyzed using one-way ANOVA followed by Fisher’s protected least significance difference [PLSD] post hoc test. Significance level was accepted at p < 0.05.

## 3. Results

### 3.1. Effect of increasing concentrations of reduced glutathione on cell index

Figure 1 shows the effect of increasing concentrations of glutathione on the cell index of lung epithelail cell A549. Cells showed a dose-dependent decrease of cell index in response to increasing GSH concentrations (Figure 1A). Exposure to increasing NAC concentrations also induced a dose-dependent reduction of cell index, although reduced compare to the effects of GSH (Figure 1B). The xCELLigence biosensor system allows the continuous measurement of cell index which is in fact an impedance, it depends on cell number and surface occupied by cells, and on adhesion strength of cells to the substrate. Significant decrease in cell index may sign cell death or cell adhesion reduction. To further understand the effect of glutathione, we observed whether glutathione decreased cell number. A 24 hour image acquisition showed that 10 mM glutathione did not induced cell death. Figure 2A shows micrographs obtained before and 8, 16, 24 hours after exposure to 10 mM glutathione. No cell death was observed but instead a marked cell surface reduction over time is obvious at 8, 16, 24 hours (Figures 2A). Furthermore, cell labeling with Zombie Red, a fluorescent marker of cells with permeable cell membrane related to cell death induction, confirmed that glutathione did not altered cell integrity even at high concentrations (Figure 2B).

**Figure 1:**
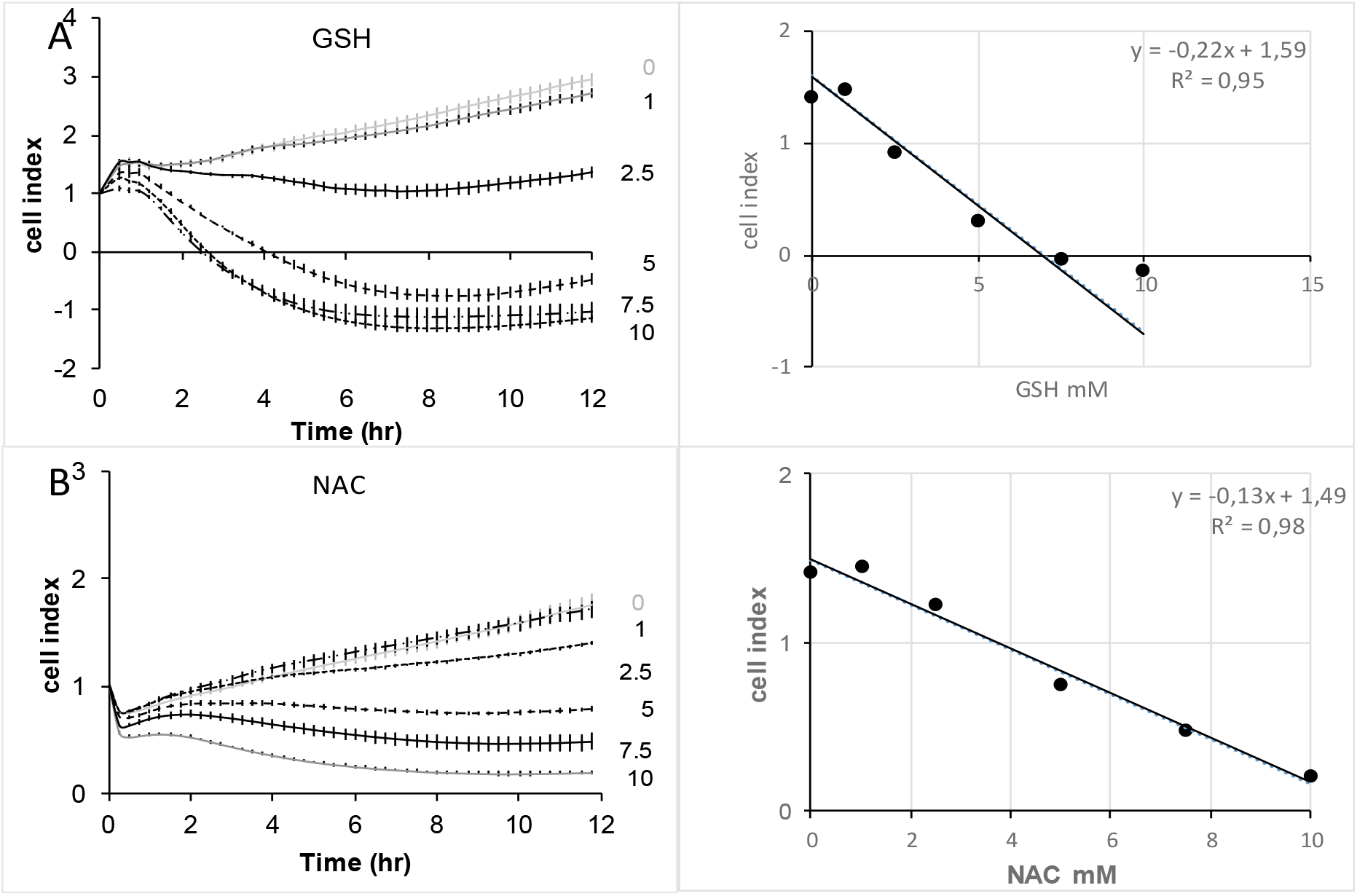
Real-time evolution of cell index in response to increasing concentrations of reduced glutathione (top) or NAC (down) on A549 pulmonary cancer cells. Cell index was normalized at the time of treatment. Data are presented as mean cell indexes ± SEM during time (left panel) and correlation curves 8 hours after treatment (right panels), n=8 replicates.

**Figure 2:**
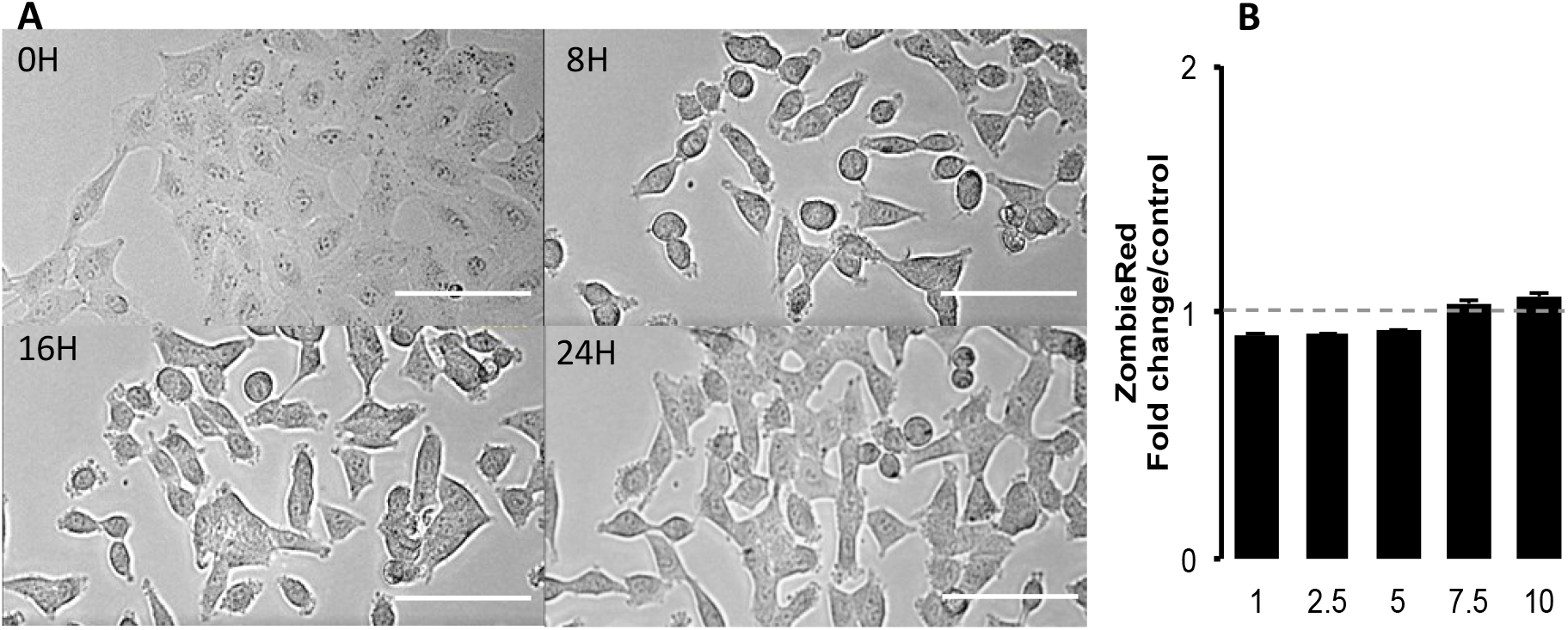
Morphological changes induced by glutathione in A549 cells. **A**: Contrast phase photomicrographs at times 0, 8h, 16h and 24h of a time-lapse in presence of 10 mM glutathione. No image of dead cell is visible. Compare to time 0, cells reduced their surface. Scale bar represents 100µm. **B**: Cell permeability to Zombie Red analyzed by fluorescence quantification in cytometry 2.5 hours after treatment, in a dose-dependent response to glutathione (mM). Data are presented as mean ± fold changes Zombie Red fluorescence intensity in reduced GSH treated cells *versus* control media (n=4).

To state whether the effect of gluthatione results from reduced or oxidized gluthatione, GSH was challenged by hydrogen peroxide. Increasing concentrations of hydrogen peroxide were incubated in the culture medium plus 10 mM GSH. The addition of hydrogen peroxide decreased the effect of GSH proportionnally to its concentration (Figure 3). Interestingly ten times more hydrogen peroxide was necessary to completely neutralize the effect of 10 mM GSH.

**Figure 3:**
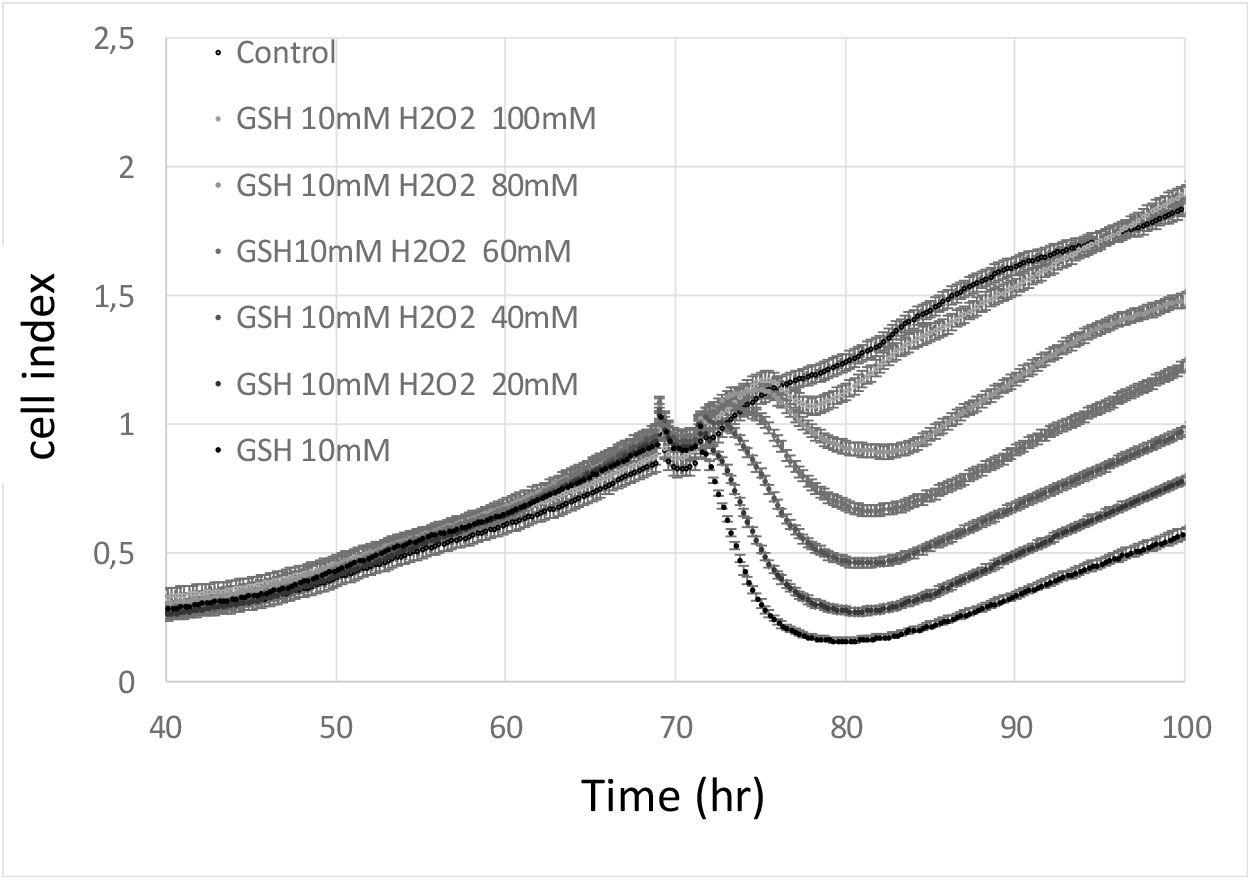
Effect of increasing concentrations of hydrogen peroxide on 10 mM GSH on cell index of A549 cells. Addition of hydrogen peroxide in the culture medium antagonized the effect of GSH on cell index in a dose-dependent manner. Results are presented as mean ± SEM (n=8).

In an other set of experiments, we investigated whether oxidized glutathione could exert some effects on cell index.

### 3.3. Effects of increasing concentrations of glutathione on cell volume

To further quantify cellular changes observed in response to GSH, we used the label-free single-cell analysis microscope Holomonitor (Phase Holographic Imaging, PHI, Lund, Sweden). The average optical volume of cells significantly increased (Figure 5A). That increase was dose-dependent and was already observable in response to 1 mM of GSH. Similar results were obtained in response to NAC (Figure 5B).

**Figure 4:**
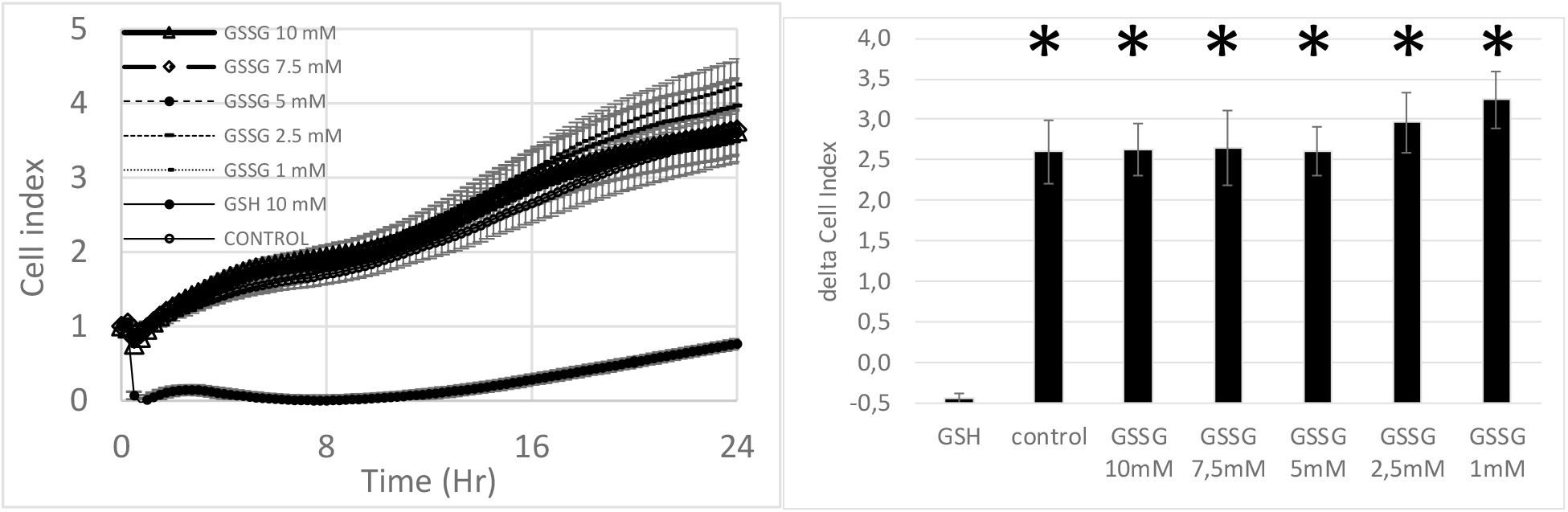
Effect increasing concentrations of oxidized glutathione on cell index of A549 cells (left panel). Addition of oxidized glutathione did not decrease cell index of A549 cellscompare to control. Results are presented as mean ± SEM (n=8). Star indicate asignificant differnce compare to GSH at p<0.05.

**Figure 5:**
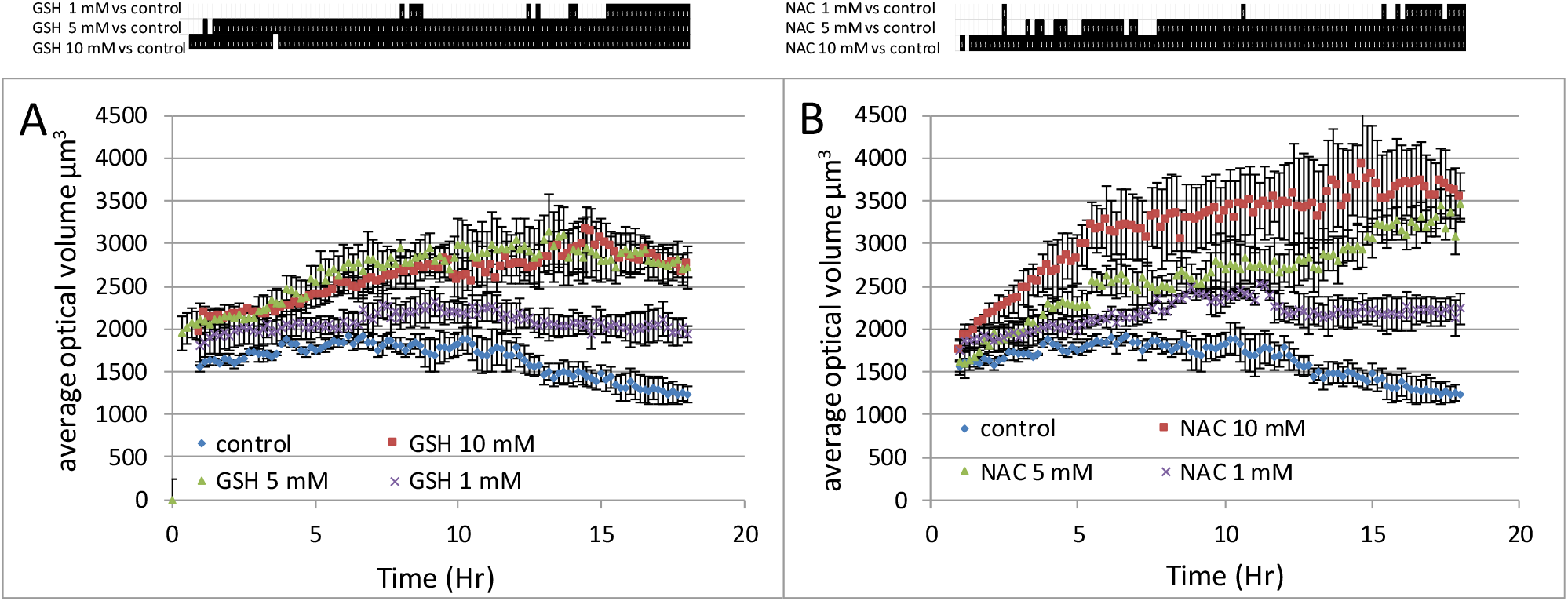
Real-time analysis of A549 cell size modifications induced by GSH (A) or NAC (B). Three different concentrations have been tested 1, 5 and 10 mM. Proliferative 459 cells were monitored on Holomonitor microscope during 18 hours. Average optical thickness of cells were registered every 10 min. Data are presented as mean values ± SEM of 4 wells with 20 cells/well. Upper part, black lines indicate significative difference compare to control at p<0.05 following one-way ANOVA analysis and Fisher’s protected least significance difference post-hoc test.

### 3.4 Inhibition of intracellular synthesis of reduced glutathione does not impair its extracellular action

In the next experiment, buthionine sulfoximine (BSO, 10mM), the inhibitor of γ-glutamylcysteine synthetase was added during 24 hours to A549 cell culture (Figure 6). The inhibitor alone did not modulate cell proliferation. It had no effect on GSH action since the same inhibitory effect was observed in response to 10 mM in presence as well as in absence of BSO. Indeed, cell index reached values close to zero after GSH addition independently of BSO. Nevertheless cell index did not reached zero indicating that cell were not dead. The fact that the inhibitor of γ-glutamylcysteine synthetase did not alter the response of glutathione is of importance, since it demonstrates that the increase of intracellular GSH concentration is not required to reduce the cell index. This is confirmed by the measure of intracellular concentration of GSH (Figure 9). BSO markedly decreased intracellular GSH concentration which did not hampered its effect on cell index.

**Figure 6:**
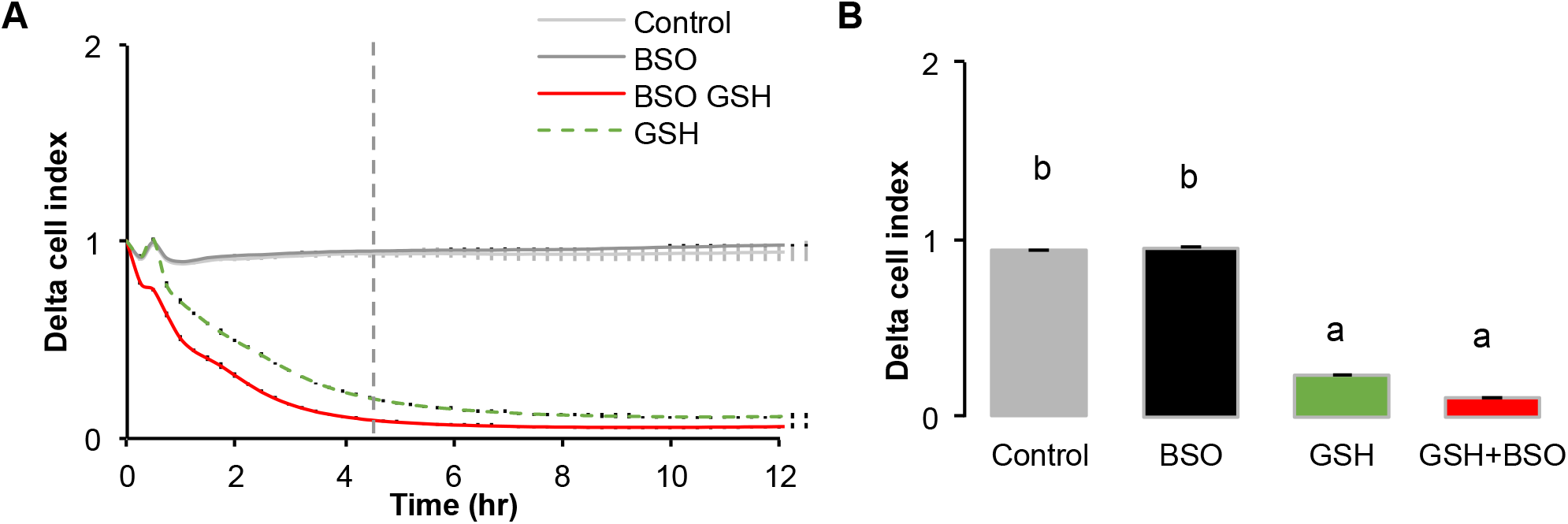
Realtime analysis of the effect of glutathione (10 mM) in presence of γ-glutamylcysteine synthetase specific inhibitor, buthionine sulfoximine (BSO) on 549 cells. A549 cells were pre-incubated with BSO (10 mM) during 24 hr. Then GSH 10 mM was added either alone or in presence of BSO. A: the graph shows a similar response between cells pre-incubated in presence of BSO or not, suggesting that the decrease in cell index is not dependent of the intracellular concentration of GSH. Similarly, BSO had no effect on control. B: histogram of delta cell indexes of control cells or cells exposed to BSO alone or GSH + BSO. Data are presented as mean delta cell indexes +/-SEM (n=8). Different letters on the top of the columns indicate a statistical significant difference after one-way ANOVA followed by Fisher’s protected least significance difference at p < 0.05.

Pro-GA is a pro-drug type of N-glutaryl-L-alanine, which penetrates into cells and inhibits the γ-glutamylcyclotransferase (GGCT) which catalyzes the chemical reaction (5-L-glutamyl)-L-amino acid to produce 5-oxoproline and L-amino acid. 5-oxoproline is a precursor of glutamate. The inhibition of GGCT decreases the intracellular production of GSH. Results show that the ProGA has no effect of cell index either in control or on the effect of GSH or NAC (Figure 7) on cell index.

**Figure 7:**
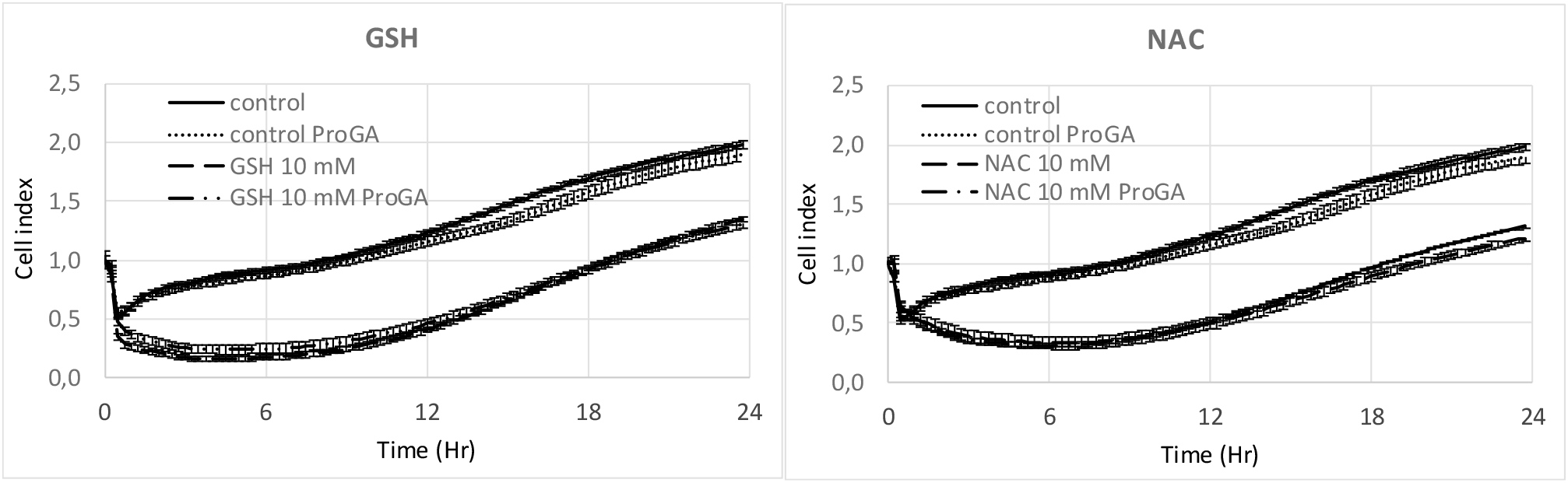
Effect of ProGA, a γ-glutamylcyclotransferase inhibitor (GCCT) involved in glutathione metabolic cycle. ProGA (100 µM) had no effect on control and GSH or NAC effects.

### 3.5. Effects of increasing GSH concentrations on cell adhesion

Since the previous experiment argued for an extracellular effect of reduced glutathione, we next tested whether it may affect cell adhesion. Cell adhesion monitoring on xCELLigence has been previously demonstrated and can be assessed in short time experiments [13, 14]. After trypsination, increasing concentrations of GSH were added to cell suspension in DMEM. Then cell adhesion was measured on xCELLigence biosensor. Results show an inverse correlation between reduced glutathione concentration and cell adhesion, the highest concentrations of glutathione corresponding to the lowest cell adhesion (Figure 8).

**Figure 8:**
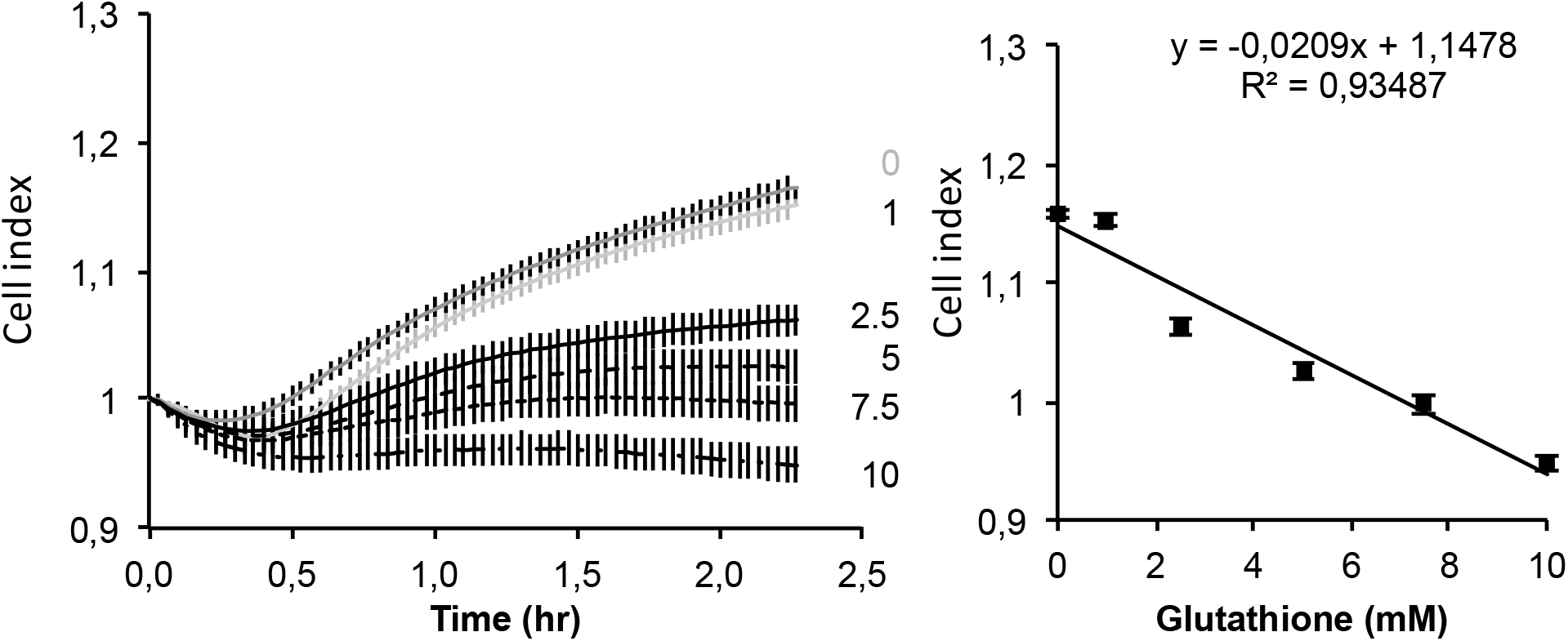
Realtime analysis of GSH effect on A549 cell adhesion. A549 cells were loaded at 5000 cells/cm^2^. Results are presented as mean delta cell indexes ± SEM (normalization to time of loading) during 2.5 hours (left panel). Correletion curve presented in right panel indicates a strong corelation between GSH concentrations and the decrease in adhesion measured by the cell index (n=8).

### 3.6. Effects of increasing concentrations of glutathione on cell cytoskeleton

The decreased cell adhesion by GSH may reflect intracellular modifications such as actin polymerization. To test that hypothesis we used phalloïdin, the toxin found in cap mushroom, known to bind specifically at the interface of F-actin subunits. FACS analysis of A549 cells labeled with fluorescent TRITC-phalloïdin shows a decreased incorporation of fluorescently labeled molecule following 24 hour GSH treatment in A549 cells attesting a decrease in F-actin (Figure 9).

**Figure 9:**
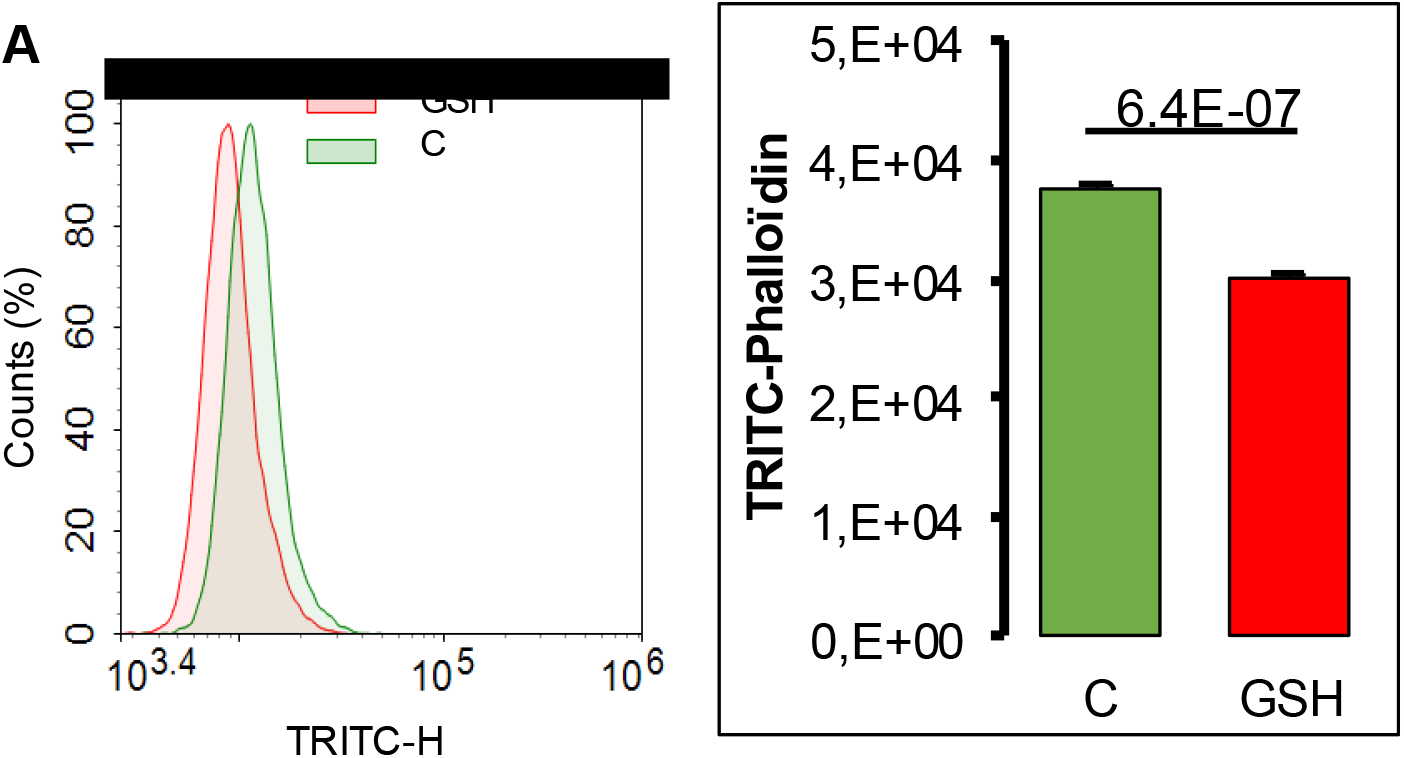
Quantification of TRITC-phalloïdin by cytometry in A549 cells exposed to glutathione (GSH 10 mM) during 24 hours. A-Representative histogram of cell number repartition according to fluorescence incorporation. B-TRITC-phalloïdin fluorescence quantification according to treatment (mean values +/SEM, n=4) with significant Student t-test p-value (p<0.05).

### 3.7. Effects of exposure to increasing GSH concentrations on oxidative stress and its intracellular concentration

Exposure of A549 cells to 10 mM GSH for 24 hours reduced oxidative stress from 2.5 to 10 mM, compare to control cells (Figure 10).

**Figure 10:**
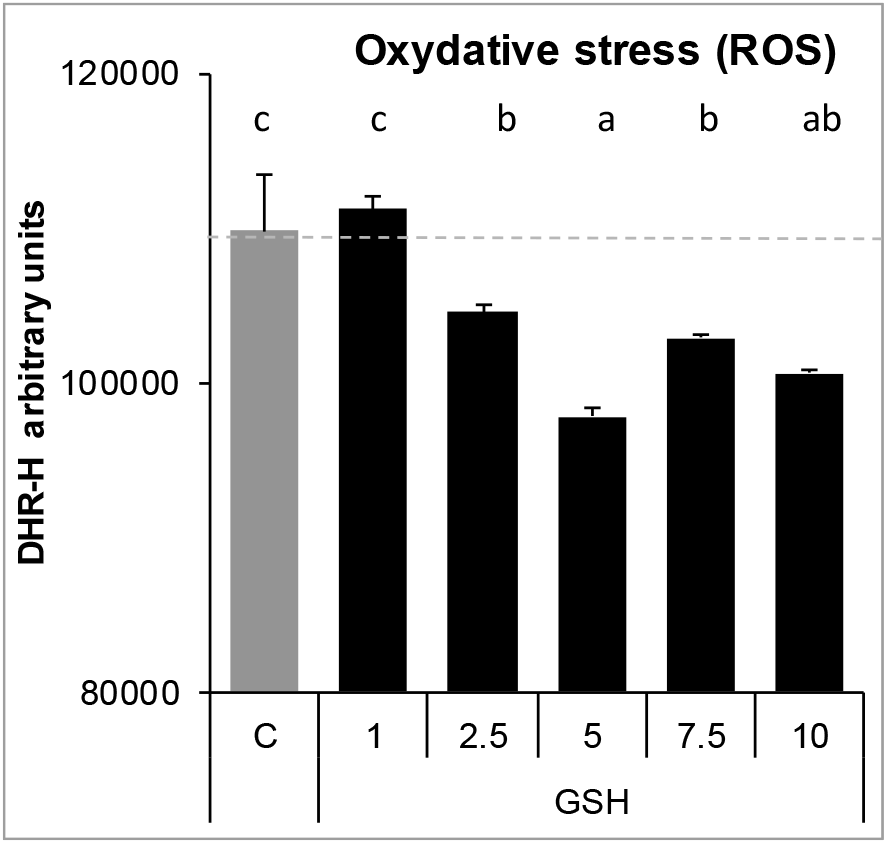
Effects of increasing GSH concentrations on oxidative stress measured with DHR (Dihydrorhodamine 123) and analyzed by FACS. Results are presented as mean± SEM. Different letters indicate significative differences after one way ANOVA followed by Fisher PLSD at p<0.05.

Exposure of A549 cells to either GSH or NAC (10 mM) significantly increased intracellular GSH concentration. Exposure of cells during 24 hours to BSO (10 mM) strongly decreased the intracellular GSH concentration despite the exposure of cell to GSH or NAC during 24 hours (Figure 11).

**Figure 11:**
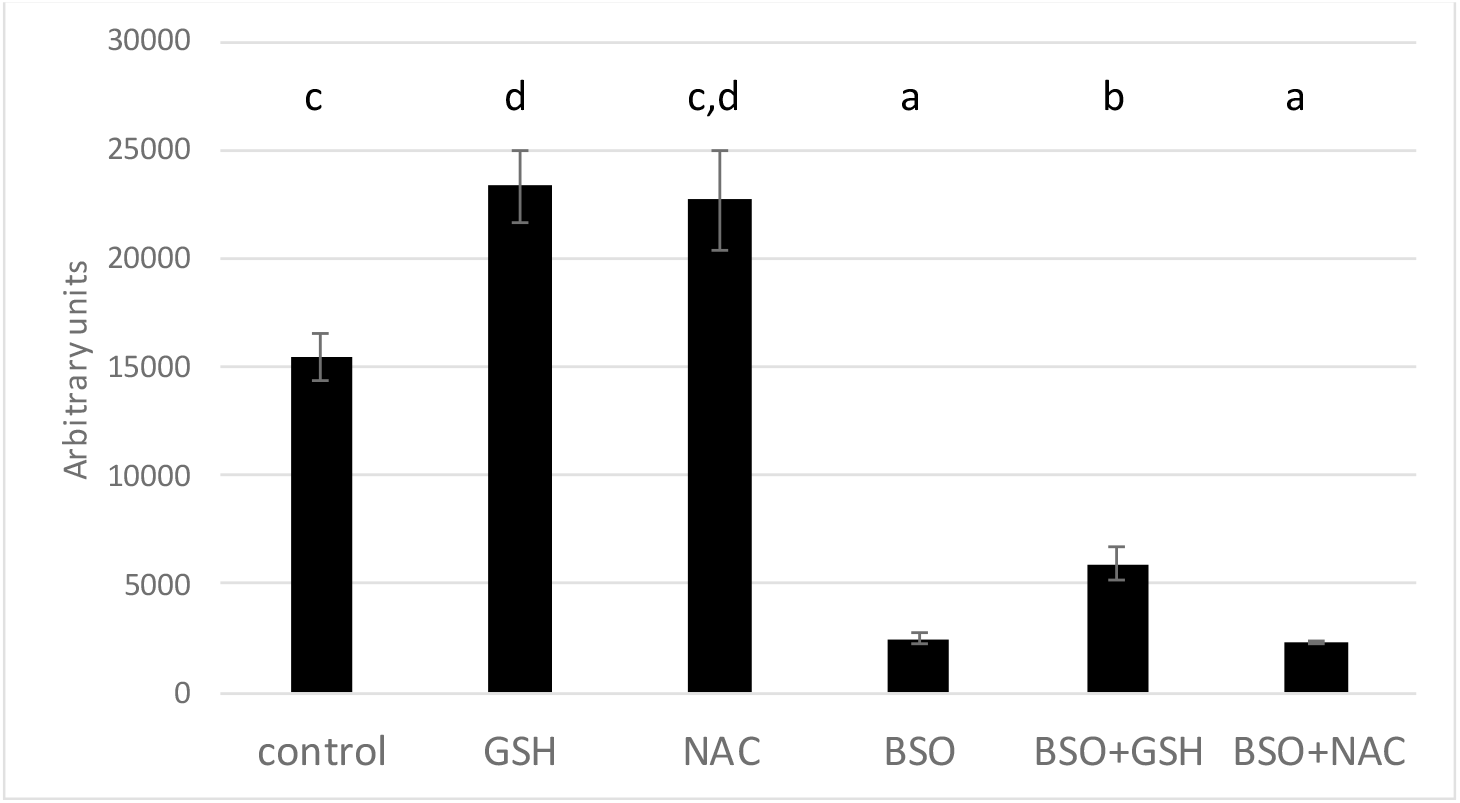
Intracellular GSH concentration after 24 hours exposure of A549 cells to GSH or NAC (10 mM) or 24 hours to BSO (10mM) and then 24 hours to BSO+GSH or BSO+NAC. Results are presented as mean ± SEM. Different letters indicate significative differences after one way ANOVA followed by Fisher PLSD at p<0.05.

## 4. Discussion

The present results show a marked effect of GSH on cell volume. Following the addition of GSH in the culture medium, the decrease in cell index is fast and marked reaching almost basal values, occurring within 2-3 hours (Figure 1A). The decrease in cell index is dose-dependent to GSH concentration (Figure 1). Similar effects are observed in response to NAC, although reduced in amplitude compare to GSH (Figure 1B). GSH markedly reduced cell index, the visual control using microscopy certified the absence of dead cell (Figure 2A). Furthermore, cells maintained their membrane integrity, attested by Zombie red labeling (Figure 2B). Finally, the decrease in cell index is reversible. Indeed, after longer time exposure, cell indexes slowly increase despite the presence of reduced glutathione (Figuer 3, 4 and 7). After washing out, the cell index increased markedly without fully reaching the control values (not shown). On xCELLigence, the cell index depends on the surface occupied by cells but also on cell adhesion strength. Since cell number is constant the only one explanation for the decreased cell index is that cell adhesion is affected by GSH. Real-time analysis using the quantitative phase contrast microscope Holomonitor M4, confirmed that GSH did not kill cells but resulted in a marked increase in average optical cell volume (Figure 5). That increase is dose-dependent. A designed experiment on xCELLigence biosensor dedicated to the measure of cell adhesion shows a dose-response to GSH on A549 cell adhesion, thus strongly arguing for a diminished cell adhesion force induced by GSH, which was confirmed by a reduction of F-actin required for integrin clustering needed for cell adhesion (review in [15]).

Whether GSH regulates cell adhesion through extracellular action rather than increased intracellular contents was supported by the extremely fast cell adhesion force reduction observed in real-time experiments. Indeed, a significant effect was detected within few minutes even at low concentrations after GSH addition. In order to test a possible effect of intracellular glutathione synthesis, buthionine sulfoximine (BSO), a specific inhibitor of γ-glutamylcysteine synthetase, was used to block the first step of intracellular glutathione synthesis. BSO at 10 mM significantly decreased intracellular GSH concentration, even after incubation with 10 mM gluthatione (Figure 11). BSO alone did not significantly change the cell index of A549 cells (Figure 6). Pre-incubation of A549 during 24 hours with BSO before GSH addition, did not change the decrease in cell index produced by GSH alone, despite the very low intracellular GSH concentration (Figure 11). It implies that GSH exerts its reducing effect on cell adhesion independently of its intracellular concentration. A second inhibitor was used to state the effect of intracellular GSH. Pro-GA inhibits the γ-glutamylcyclotransferase (GGCT) which produces 5-oxoproline a glutamate precursor. Here again, the inhibition of intracellular GSH production had no impact on the effect of GSH on cell index, arguing for an extracelullar action (Figure 7A). Similar results are observed with NAC (Figure 7B). Since GSH can be easily oxidized, the effect of oxidized glutathione was also tested on cell index. Results are clear, oxidized glutathione did not decrease cell index. The reduction of cell index is triggered by GSH (Figure 4). In an other set of experiments, GSH was oxidized by increasing concentation of hydrogen peroxide (Figure 3). A perfect dose-response was obtained on cell index by GSH. It is noticeable that ten times more hydrogen peroxide was necessary to fully antagonize the effect of GSH. This may lie from the fact that hydrogen peroxide is highly reactive and not specific to GSH.

Concerning the mechanism involved in the action of GSH on cells, data from the literature show that both GSH and N-acetylcysteine (NAC) are disulphide breaking agent [16, 17]. NAC produces similar effects as GSH on cell index, average optical volume (Figure 1 and 5) and phalloïdin labeling (results not shown). Disulfide bonds formed by the oxidation of two cysteine residues to give cystine are essential for the stability of secreted and plasma-membrane proteins (reviewed in [18]). Their cleavage is also of most importance to regulate secreted proteins such as thrombospondin, an extracellular glycocoprotein involved in cell-matrix adhesion. The role of disulfide bounds in the stability of focal adhesion proteins are largely depicted in the control of cell surface receptors [19] as well as the role of extracellular protein disulfide isomerases in the regulation of integrins, in platelet adhesion and thrombosis [10] and more recently in cancer cell migration and invasiveness [8, 20]. In breast cancer cells, the regulation of disulfide exchanges, mostly involving integrins, controls transendothelial migration of cancer cells. Such mechanism has been demonstrated using several free thiols blocking agents and/or protein disulfite isomerase inhibitors.

The effect of reduced glutathione rises numerous questions. The GSH concentrations used in the present study are high. Some experiments have been performed with 10 mM but several experiments have tested dose-responses effects from 1 to 10 mM. Significant effects of GSH are observable at much lower concentrations than 10 mM. It is the case for xCELLigence cell index which is a highly sensitive method, allowing to detect and quantify cell adhesion modifications in shorter times (few minutes) than fluorescence imaging or cytometry in TRITC-phalloïdin labeled cells (2.5H). Several studies have reported rather high intracellular GSH concentrations, for example in liver cells it reaches 10 mM [2]. Plasma and extracellular pools of GSH are much lower than the intracellular pool, around 1-10 µM at the exception of bile 10 mM and alveolar lining fluid, 400-800 µM [2, 21, 22, 23]. Present results also rise question on the true extracellular GSH concentration. Indeed, if GSH combines with disulfide bridges on extracellular proteins, it then becomes obvious that its extracellular concentration drops, but then the extracellular GSH concentration takes into account only free GSH, which cannot be used as a reference for GSH excreted by cells.

Export of GSH out of cells was observed in several cell types: human lymphoid cells, skin fibroblasts and macrophages suggesting that it is a commune feature. It may reduce compounds in the immediate environment of the cell and might protect cells [23]. Present results demonstrate that it can also control cell adhesion. Cell adhesion is mediated through integrins, a family of α/β heterodimeric molecules responsible for cell–matrix and cell–cell interactions. Integrins are subject to redox dependent conformational changes resulting in alterations of their binding activity [6]. All integrins contain a large number of cysteine residues in their ectodomains. The breakage of disulfide bonds in integrins by reducing agents can induce their activation [10, 24, 25]. Integrin activation by thiol-disulfide was described on αVβ3, αIIbβ3, integrin subunit α11 and β1 integrin [20]. Furthermore, the protein disulfide isomerase (PDI) directly interacts with integrins, mediating disulfide-thiol exchange, resulting in conformational changes and activation. PDI can be found on the surface of several cell types, including fibroblasts [20]. Adhesion of human skin fibroblasts to collagen and fibronectin is inhibited by para-chloro-mercuriphenyl sulfonic acid (pCMPS), a thiol blockers [4, 7].

## 5. Conclusions

The present study demonstrates that GSH has a major impact on cell adhesion. That decrease in cell adhesion results in an increased cell volume. That rises the hypothesis that GSH produced by cells can exert a direct control on cell adhesion and cell volume. Such a hypothesis adds a new function of glutathione and most likely will necessitate a re-evaluation of cell adhesion functions according to their redox-state.

## Author contributions

(Conceptualization, A.G.; methodology, A.G. and E.B.; validation, A.G. and E.B., investigation, A.G. and E.B.; data curation, A.G. and E.B.; writing—original draft preparation, A.G. and E.B.; writing—review and editing, A.G.; supervision, A.G.; project administration, A.G.

